# Assessment of Structural Units Deletions in the Archaeal Oligosaccharyltransferase AglB

**DOI:** 10.1101/2020.08.18.256065

**Authors:** Conrado Pedebos, Hugo Verli

## Abstract

Oligosaccharyltransferases (OSTs) are enzymes that catalyze the transfer of a glycan chain to an acceptor protein. Their structure is composed by a transmembrane domain and a periplasmic / C-terminal domain, which can be divided into structural units. The *Archaeoglobus fulgidus* OST, AfAglB, has unique structural units with unknown functions. Here, we evaluate the stability role proposed for AfAglB units by employing molecular modelling and molecular dynamics simulations, to examine the effect of single and double deletions in the enzyme structure. Our results show a strong effect on the dynamics of the C-terminal domain for the mutated systems with increased fluctuations near the deleted areas. Conformational profile and stability are deeply affected, mainly in the double unit deletion, modifying the enzyme behavior and binding interfaces. Coordination at the catalytic site was not disrupted, indicating that the mutated enzymes could retain activity at some level. Hotspots of variation were identified and rationalized with previous data. Our data shows that structural units may provide stabilization interactions, contributing for integrity of the wild-type enzyme at high temperatures. By correlating our findings to structural units mutagenesis experimental data available, it was observed that structural units deletion can interfere with OSTs stability and dynamics but it is not directly related to catalysis. Instead, they may influence the OST structural integrity, and, potentially, thermostability. This work offers a basis for future experiments involving OSTs structural and functional characterization, as well as for protein engineering.

## Introduction

N-glycosylation is a co-/post-translational modification widespread among all domains of life. This process is involved in many actions and activities throughout cells, such as stability, signalling, and immune response in *Eukarya*^1^; flagella assembly, motility, protection against extreme environments, and integrity of structures in *Archaea*^2^; host adhesion, invasion, and colonization in *Bacteria*^3,4^. Despite the intrinsic differences for each species, the overall N-glycosylation pathway share many similarities^5^. It is comprised by many enzymes that sequentially add monosaccharides to a lipid carrier which is, then, translocated by a flippase to the exterior side of the cell (in *Archaea*), to the periplasmic space (in *Bacteria*), or to the lumenal face of the ER (in *Eukarya*). In *Archaea*, the final step of the pathway is catalyzed by the oligosaccharyltransferase AglB, an enzyme that transfers the lipid-linked oligosaccharide (LLO) to an acceptor protein carrying the sequon N-X-S/T (that is, Asn-any amino acid residue except Pro-Ser or Thr)^6^.

Recently, the full structure of two prokaryotic OSTs were elucidated: PglB from *Campylobacter lari* (ClPglB)^7,8^, and the longest paralog AglB from *Archaeoglobus fulgidus* (AfAglB-L) (Figure 1)^9^. Before that, only partial structures for the C-terminal globular domains were obtained^10–14^, demonstrating that the archaeal domain of life possesses the greatest variability. As seen in these previous works, the structure of an OST can be divided in two large domains: a transmembrane domain (TM) consisting of 13 α-helices, while also having external loops (EL) that display important catalytic residues^7^, such as EL1, EL2, and EL5; and a C-terminal globular domain, which usually consists of an α/β mixed fold. This domain can be further subdivided into structural units. Firstly, the central core (CC) unit is the only substructure that is essential to every OST. It participates in the interface region with the TM domain, assembling the enzyme in two binding sites (protein acceptor site and LLO donor site). Accordingly, this unit holds the WWDYGY (WWDXGX) conserved motif^10,15^, responsible for the recognition of the S/T (+2) of the N-glycosylation sequon, and the DK motif^7,10^, which keeps the (+2) position residue in a tight binding by providing additional interactions. The second unit is the Insertion Sequence (IS), which receives this name because it seems to be inserted in the CC unit area^14^. Generally, it displays an organization of β-strands. For example, ClPglB and *P. furiosus* AglB (PfAglB) possess a distorted β-barrel fold, while AfAglB-L presents a reduced unit without the β-barrel organization. As a substituent for the barrel structure, a three-helices bundle (TH-Bundle) (originally classified as being part of the CC unit)^14^ is located at the same place and seems to play that role, a unique feature in the OSTs structures found until now, including the partial structures. The two final structural units are the Peripheries 1 (P1) and 2 (P2), only identified in Archaeal structures. These units are β-sheet rich, with slightly different organizations, and they are distributed around the CC unit, on the opposite direction of the IS unit. Archaeal OSTs can display either only P1 or both units at the same time, but no structure carrying only P2 was determined yet.

**Figure 1.**
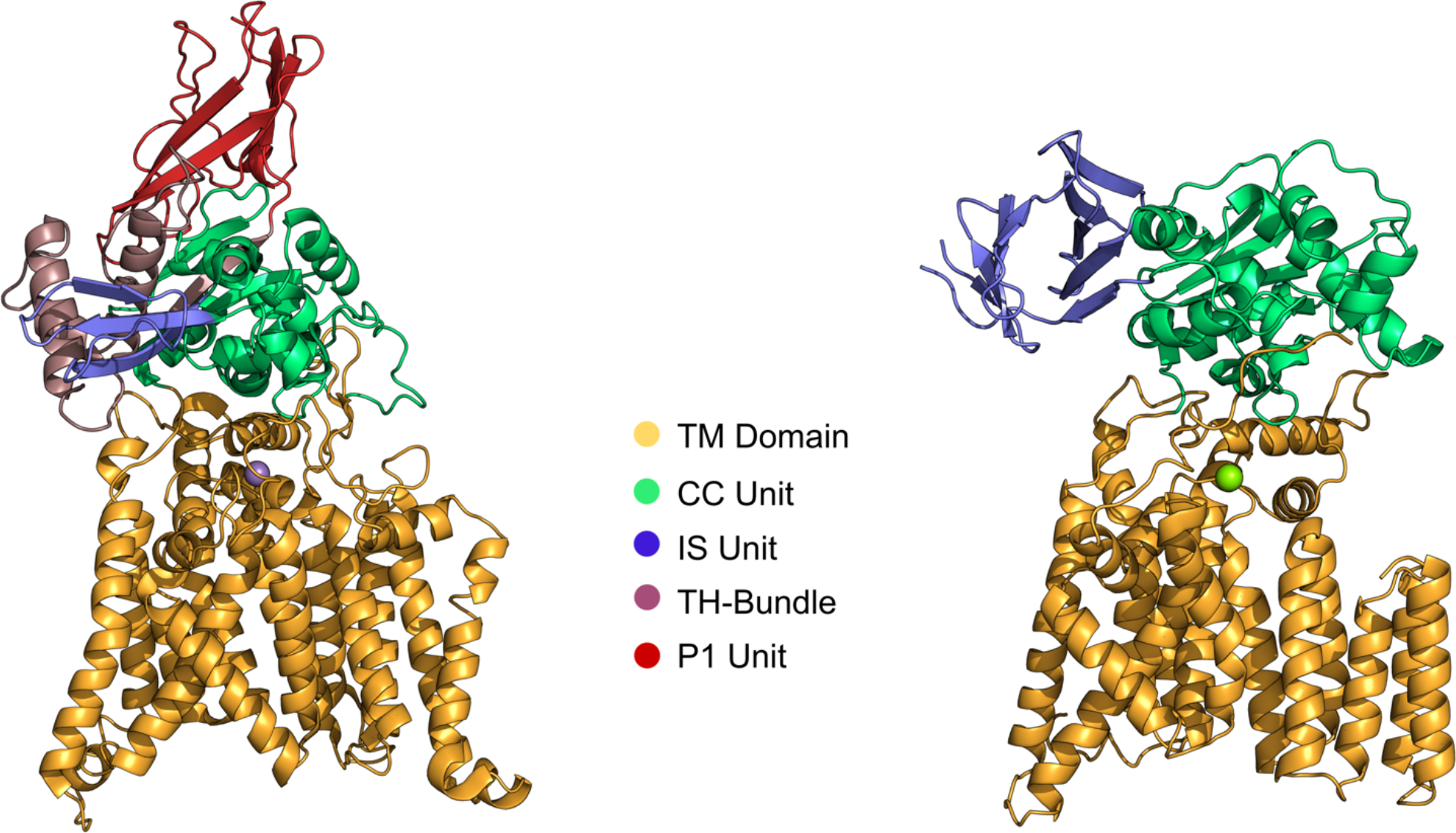
AfAglB-L (*left - PDB ID: 3WAK*) and ClPglB (*right - PDB ID: 3RCE*) crystallographic structures. The TM domain (orange) and the C-terminal domain structural units (CC-green, IS - blue, P1-red) are depicted. The purple sphere corresponds to Zn^2+^ ion, while the green sphere corresponds to Mn^2+^ ion. The TH-Bundle is originally part of CC unit, although it occupies the same place of the IS unit from ClPglB and interacts with the IS unit from AfAglB.

Despite all of this information, until today only one experimental study assessed structural units properties^11^. In this previous work by Matsumoto and co-workers, a mutant PfAglB with a deleted IS unit (PfAglBΔIS) displayed the same activity than the wild type enzyme, indicating that this unit is not mandatory for catalysis. However, the assays were performed with detergent-solubilized PfAglB, in a temperature lower than optimal, and a short peptide as an acceptor, caveats that could interfere with proper evaluation of the enzyme function^11^. Two hypotheses emerged for the functions of the IS unit: structural stability when the protein is embedded in a membrane, and support for the regulation or efficiency of the enzyme’s activity when in the presence of larger polypeptide chains and when coupled to protein translation processes^11^. As for P1 and P2, the only current hypothesis^14^ is that they could contribute for the thermostabilization of the enzyme under extreme conditions, since the extra units (and specifically P1 and P2) are only found in this domain of life. Another mentioned possibility is that these units could be equivalent to the subunits that constitute the OST multimeric complex from eukaryotic species (5-8 subunits). Recent structural characterizations of these eukaryotic OST complexes^16,17^ have demonstrated the presence of a β-sheet rich subunit (Wbp1) in areas similar to the ones occupied by P1 and P2 units in archaeal OSTs (Figure S1). However, this subunit presents distinct folding and organization compared to P1 and P2, along with a transmembrane portion not observed in those units. Although they might play similar roles in the process, the lack of similarity and the still unknown function of Wbp1 prevent us to make further inferences.

Investigations of OSTs mechanisms could lead to the production of tailor-made glycoproteins. In fact, recent data have demonstrated modifications enhancing OSTs catalysis, such as mutations of residues to increase the recognition and binding of acceptors^18-20^, change of the catalytic ion to increase the rate of product formation^9,21,22^, as well as the production of different sizes of lipid carrier and interspecies glycan chains^23,24^. However, there is still a lack of studies evaluating the role of IS, P1, and P2 structural units, which could bring new insights into the enzyme functionality. Therefore, in this work, we employed molecular dynamics (MD) simulations to observe the structural effects caused by full deletions of AfAglB-L extra structural units. Specifically, we aimed to: I) evaluate the mutants stability after the deletion of P1, and P1 plus IS units; II) verify the behavior of the catalytic site in order to analyze if the requirements for catalysis are maintained upon deletion of the domains; III) identify important interactions between structural units that may govern the protein stability under hyperthermophilic conditions, allowing inferences regarding these units transferability between distinct OSTs. Our work seeks to contribute for the comprehension of the structural basis that governs OSTs dynamical behavior, providing insights for potential biotechnological applications of these enzymes.

## Materials and methods

### Systems Preparation

For this study, the structure from the RCSB Protein Data Bank under the PDB ID: 3WAK^9^ was retrieved, since it corresponds to a full apo structure containing EL5. Mutants were obtained by deleting the P1 region (residues 777-868), and the IS region plus the TH-Bundle (residues 638-776), including a short region from the CC unit that connects IS to P1. Three systems were studied in this work: i) wild type AfAglB-L in its apo state (AfAglB-L-WT); ii) AfAglB-L with the deletion of P1 unit (AfAglB-L-ΔP1); iii) AfAglB-L with deletion of IS and P1 (AfAglB-L-ΔISP1). In this work, we established that the ΔISP1 included the TH-Bundle from the CC unit. The reason for that is because it makes close contacts with the IS unit and forms a structure separated from the enzyme core, resembling the common β-barrel IS seen in other OSTs. To better describe the environment where these proteins are found, they were embedded in a 1-palmitoyl-2-oleoyl-sn-glycero-3-phosphoethanolamine (POPE) membrane with 512 phospholipids. The methodology utilized for protein insertion was based on the tools LAMBADA and InflateGRO2^25^.

### Molecular Dynamics Simulations

The triclinic simulation boxes received SPC water solvation^26^, in the presence of periodic boundary conditions. In the solvation step, Van der Waals radii of C atoms were raised from 0.150 to 0.375 to avoid water molecules filling spaces between phospholipids and protein. This parameter was returned to the default value after this step. PME^27^ was chosen for the electrostatic treatment of the system. The LINCS method^28^ was applied to constrain covalent bonds, allowing an integration step of 2 fs. GROMACS simulation suite^29^ with GROMOS54A7 force field^30^ was used for the molecular dynamics (MD) simulations. POPE membranes were described by GROMOS53A6 parameters (compatible with GROMOS54A7) following a previous report^31^. Cl^-^ counter ions were added to neutralize the residual charge of the systems, when needed.

Firstly, energy minimizations were performed using the Steepest Descent algorithm, stopping when force on the system was below 100 kJ·mol^-1^·nm^-1^ as a convergence criterion. Then, MD simulations were performed in two phases: an equilibration and a production phase. For the equilibration phase, the systems were simulated for 1 ns in a canonical ensemble (NVT) and for 20 ns in an isothermal-isobaric ensemble (NPT). Parrinello-Rahman barostat^32,33^ was applied with a coupling constant of 2.0 ps. As thermostats, V-rescale (NVT step)^34^ and Nosé-Hoover (NPT step - equilibration and production phases)^35,36^ were employed with coupling constants of σ = 0.1 and 1.0, respectively. Also, semiisotropic pressure was applied due to the presence of the membrane. A constant temperature of 356 K was implemented, since this is the optimal growth temperature for *A. fulgidus*^37^. During the equilibration phase, the protein was subjected to position restraints with a force constant of 1000 kJ·mol^-1^·nm^-2^. Subsequently, in the production phase, unrestrained MD simulations were performed for 300 ns for each system. Simulations were performed in triplicates (3 x 300 ns), starting with different velocities, aiming to enhance the sampling robustness of the conformational ensemble and remove low probability conformational events. Data was collected during the production phase and analyzed using GROMACS suite tools. VMD^38^ and PyMOL^39^ were used for trajectories visualization and for manipulation of structures.

## Results

### Global Structural Stability

We began by performing a global evaluation of the enzymes observing the deviation from the initial structures. The root mean square deviation (RMSD) (Figure 2A) plot for the whole structure indicates that both mutated systems demonstrate higher divergence in their structures when compared to the wild-type system. Particularly, the AfAglB-L-ΔISP1 mutated system appears to suffer a dramatic increase on its average RMSD, showing that the double deletion causes a much deeper impact in the protein structure. AfAglB-L-ΔP1 does not display such high variations, although there is a clear tendency of higher magnitude change compared to AfAglB-L-WT.

**Figure 2.**
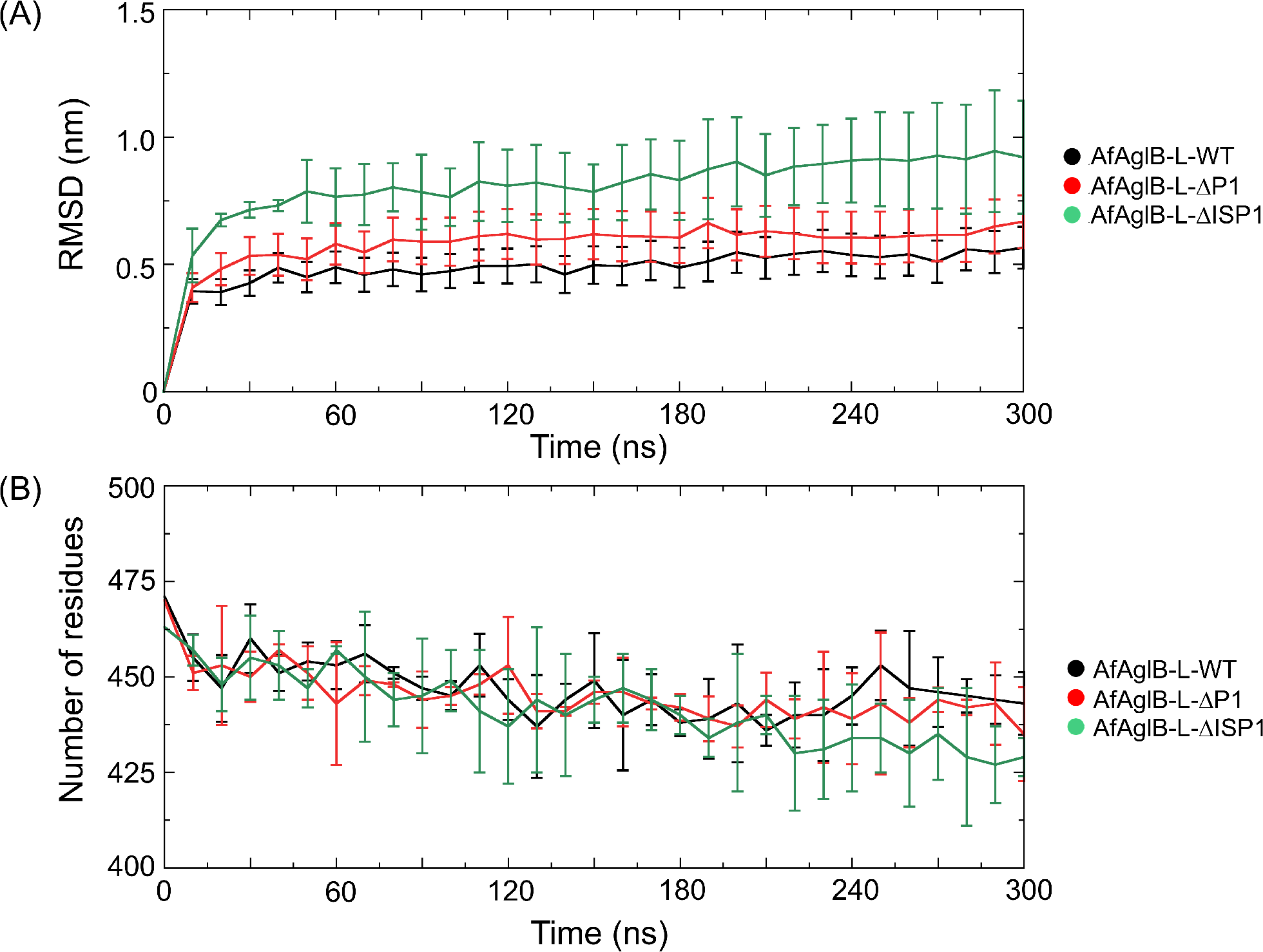
RMSD and Secondary Structure content during simulations. (A) RMSD relative to the crystal structure for each system at every 10ns with standard deviations. (B) Secondary structure content for each system at every 10ns with standard deviations. Results are shown as an average between replicates with standard deviation.

Analyzing the difference between the fluctuation of the backbone atoms as an average per residue (Figure 3), we observed that the TM domain (residues 1 to 500) maintain a generally stable behavior in its helices for all systems, with some minor variations in flexibility for the intracellular loops. EL5 shown its well-known flexibility, depicted by the large standard deviations observed in its residues.

**Figure 3.**
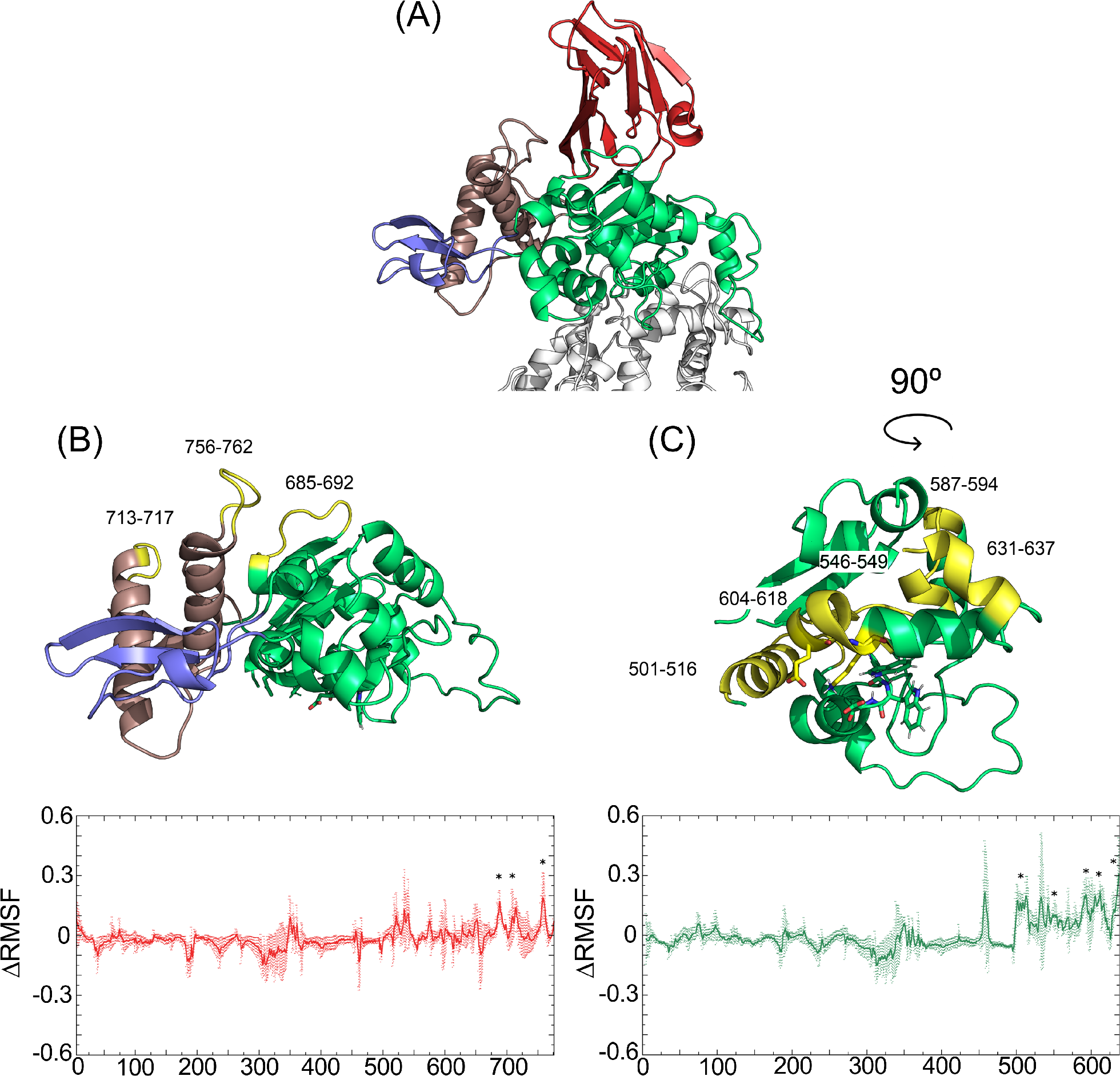
RMS fluctuation difference between backbone atoms as an average of each residue comparing the mutated systems to the wild-type. (A) AfAglB-L-WT structure of the C-terminal domain. (B) *Top*. C-terminal domain of the AfAglB-ΔP1 system: CC unit is colored in green, IS unit is colored in blue, TH-Bundle is colored in brown, and the high fluctuation areas are colored in yellow. *Bottom*. RMSF difference (red) between AfAglB-L-ΔP1 and AfAglB-L-WT with stars marking the highest fluctuations areas. (C) *Top*. C-terminal domain of the AfAglB-ΔISP1 system: CC unit is colored in green, and the high fluctuation areas are colored in yellow. *Bottom*. RMSF difference (green) between AfAglB-L-ΔISP1 and AfAglB-L-WT with stars marking the highest fluctuations areas. TM domain is hidden on (B) and (C) for better visualization. WWDXGX and DK motifs are represented in sticks.

As expected, we perceived a deeper influence from the structural units deletion in the C-terminal domain. Each mutated system displays a unique pattern of higher fluctuations when compared to AfAglB-L-WT. The single unit deletion (AfAglB-L-ΔP1) impacts areas previously interfaced with the P1 unit (Figure 3B), specially near the IS unit and TH-Bundle region: the loop (residues 713-717) connecting helices aB and aC (TH-Bundle), the loop (residues 756-762) that covalently attaches helix aA (TH-Bundle) with the core of the enzyme, and a loop (residues 685-692) that provided non-bonded interactions between the CC unit and the P1 unit. The double unit deletion (AfAglB-L-ΔISP1) deeply affected (Figure 3C) the CC unit region: the first helix of the C-terminal domain (residues 501-516), both the β-strands and the kinked-helix (residues 546-549 and residues 604-618) that carry important recognition motifs (WWDXGX and DK), as well as two other small helical areas (residues 587-594 and 631-637).

The IS unit seems to be naturally flexible, since it demonstrated some flexibility also in the AfAglB-L-WT system. However, the deletion of P1 caused an enhancement of this feature, more specifically at the TH-Bundle, which interacts with the β-hairpin structure, substituting the distorted β-barrel seen in ClPglB. To further evaluate the extent of these instabilities, we analyzed the secondary structure content of each system (Figure 2B), considering only the residues in common between all systems. All proteins had a decrease in their content of secondary structure of at least 20 residues. Both mutations appear to have a higher decrease trend on the secondary structure content, especially in AfAglB-L-ΔISP1, exhibiting the most pronounced diminishment (around 35 residues).

### Structural Units Deletion Affects the Conformational Behavior of AfAglB

To better comprehend how the mutations impact the sampling of AfAglB conformational landscape, we employed principal component analysis (PCA) on the MD simulations data. By plotting the two-dimensional subspace defined by the top two principal components (PCs) (Figure 4), we were able to evaluate the structural relaxation of the proteins individually. For the WT system, a similar behavior occurred for all replicates considering the whole protein structure, as observed by plotting the top two eigenvectors projections: an initial fluctuation, jumping among low-energy basins, as well as an apparent accommodation by the end of each simulation. Replicates 1 and 2 sample mostly one specific conformational state, while replicate 3 takes some time jumping among different regions before finding a more stable area. Despite that, the replicates explored multiple distinct areas from the subspace, indicating that our simulations possibly did not reach convergence in the simulated time. As for the mutated systems, the replicas behaviors are identical: large drifts through the conformational space, as well as no overlap between systems, indicating that these systems are even farther from convergence than the apo state simulations, which may be related to the appearance of new energy minima upon domains deletion.

**Figure 4.**
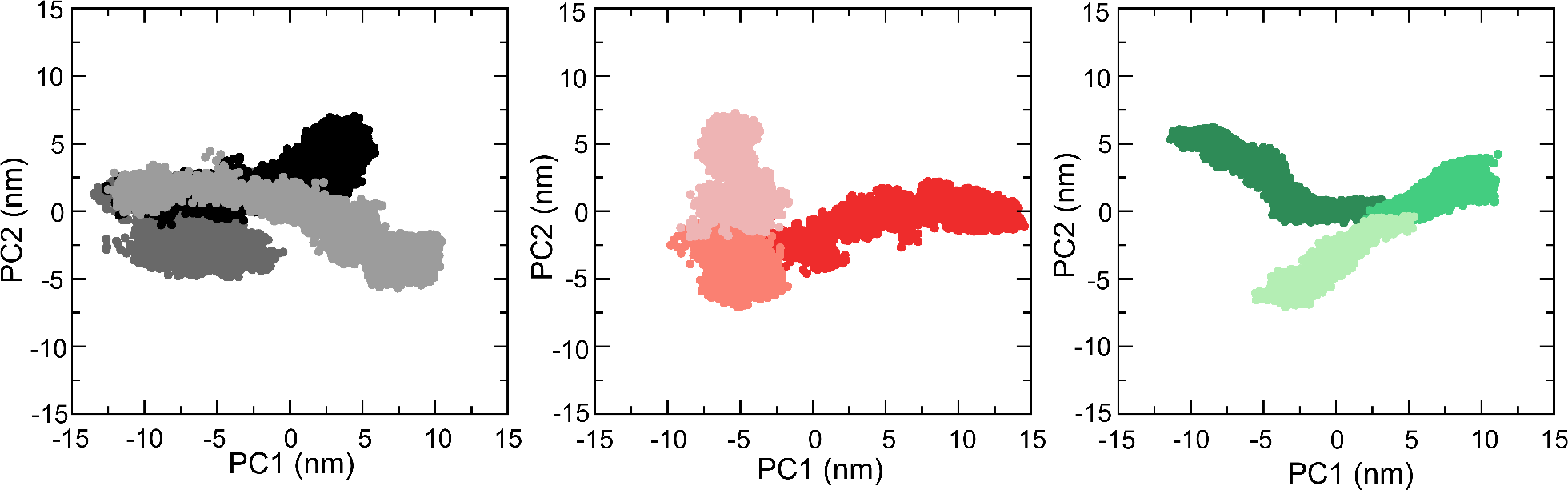
Top two PCs projections extracted with the PCA technique for the individual systems with concatenated trajectories. Individual replicates are represented by a color code of shades of black (AfAglB-L-WT), red (AfAglB-L-ΔP1), and green (AfAglB-L-ΔISP1).

To better characterize the influence of the deletions on the protein structure, we employed the same analysis for the isolated TM domain, and for the C-terminal domain structural units (CC, IS, and P1) of AfAglB-L (Figure 5). Firstly, all trajectories are concatenated into one, so that the results for different systems may be compared for a common subspace. In agreement with our observations concerning the structural divergences and fluctuations, the AfAglB-L-WT and the AfAglB-L-ΔP1 systems demonstrated a less divergent behavior, including an apparent level of overlap between some of their replicates (indicating similar conformational sampling), specifically in the TM domain and in the CC unit. In addition, these units movements are more contained at specific portions of the subspace spanned by the two PCs analyzed, a behavior which suggests that there is less conformational variation in this domain. Contrastingly, AfAglB-L-ΔISP1 is largely affected by the dual deletion, demonstrating the same drift behavior observed for the global structure. In fact, the structural adjustment suffered by AfAglB-L-ΔISP1 is so intense that subtle conformational variations from the other two systems are probably concealed.

**Figure 5.**
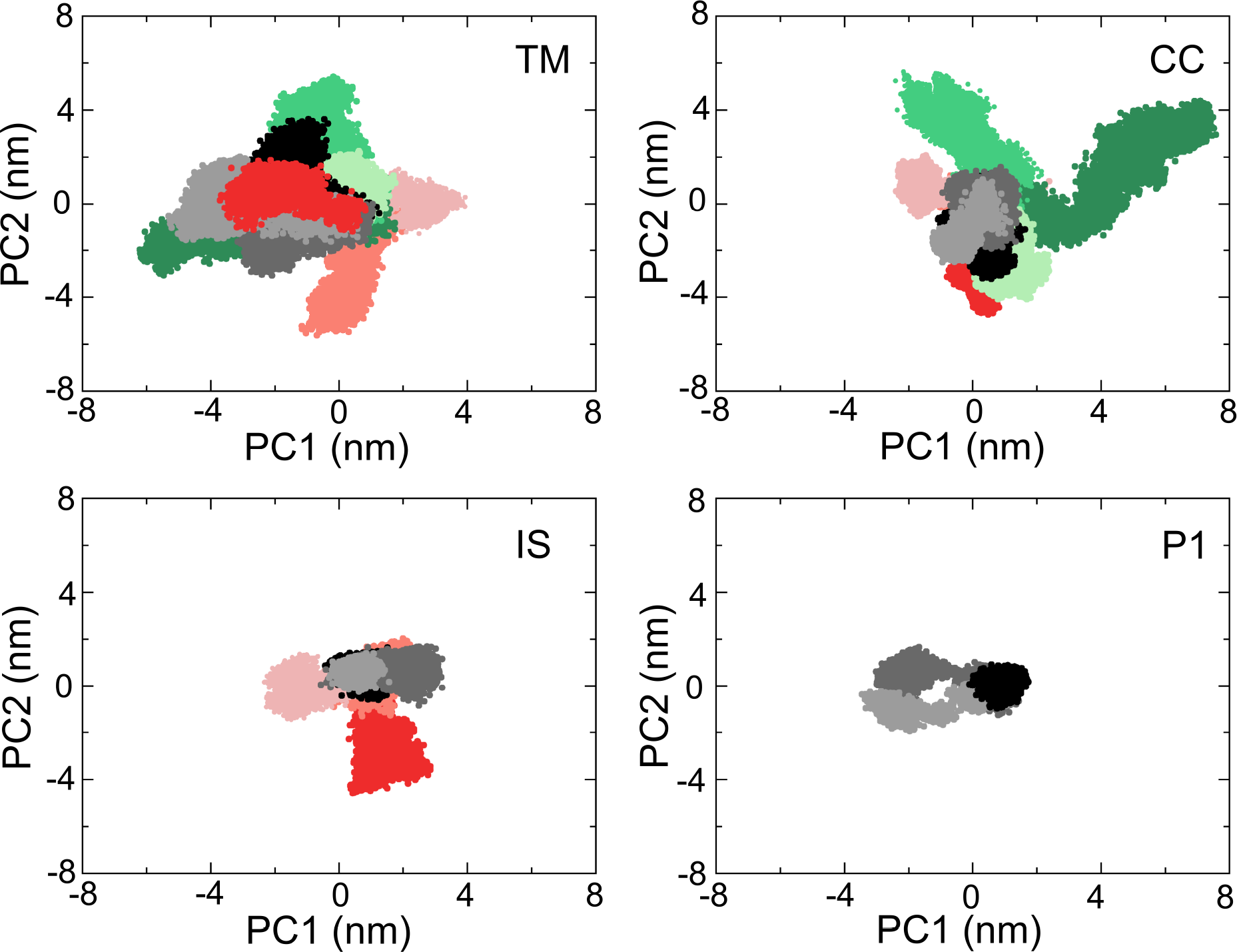
Top two PCs projections for all systems with concatenated trajectories extracted with PCA technique. Individual replicates are represented by a color code of shades of black (AfAglB-L-WT), red (AfAglB-L-ΔP1), and green (AfAglB-L-ΔISP1). Each graph represents a different domain or structural unit of the enzyme, as indicated in each legend.

The CC unit repeated the same pattern seen for the TM domain: AfAglB-L-WT and AfAglB-L-ΔP1 displayed a similar shape on the projections, while AfAglB-L-ΔISP1 maintained its dominance in the analysis with a high drifting pattern, even larger than for the TM domain, confirming that this mutation affected more the C-terminal area. Subsequently, when we analyzed the IS unit, the deletion in AfAglB-L-ΔP1 demonstrated a considerable amount of influence on this unit behavior. Replicate 2 mainly sampled the same regions as the replicate 2 of the WT system, while replicates 1 and 3 from AfAglB-L-ΔP1 drifted apart from the others, accounting for most of the variance constituting both PCs. AfAglB-L-WT replica 3 samples of the same area as replicate 1 from the same system, while replicate 2 generates a new basin. Finally, P1 unit (from AfAglB-L-WT) shown a less diffuse behavior, populating one well-defined area of the subspace where the three replicates occupied for some time during the simulations, and two other areas that were occupied separately by replicates 2 and 3, while replicate 1 stayed trapped near its initial region.

### Binding Site Coordination and Interface Stability

An important feature of OSTs, as metalloenzymes, is the maintenance of an organized network of interactions between the catalytic residues, as well as the metal ion coordination geometry. Consequently, aiming to assess the impact of the deletions on this network, we evaluated these interactions during the MD simulations by calculating the distances concerning the atoms from the side-chain of essential residues (Table 1). From that, it was possible for us to observe that all systems maintained a mostly similar coordination geometry to the one found on the crystallographic structure (PDB ID: 3WAK), although with a couple of differences: His163, which showed an increase on its distance relative to Zn^2+^, and the presence of more water molecules surrounding the metal ion (3:1), similarly to the coordination proposed on crystal form 1 in the same work by Matsumoto *et al*. (PDB ID: 3WAJ)^9^. Glu360, a presumed critical residue for the catalysis mechanism (equivalent to Glu319 in ClPglB), did not interacted with the metal ion, thus not forming the complete active site, as detected in the crystallographic structure. The metal ion remained in a tight binding at the enzyme core, with very rigid distances between the metal ligands (max. standard deviation of 0.4 Å), and one of the water molecules bridging an interaction involving His163 and Zn^2+^. We identified fluctuating coordination numbers, between 6 and 7, depending on the denticity of the carboxylates from Asp and the number of water molecules involved (Figure 6), exhibiting either a distorted octahedral shape or a face-capped octahedron.

**Figure 6.**
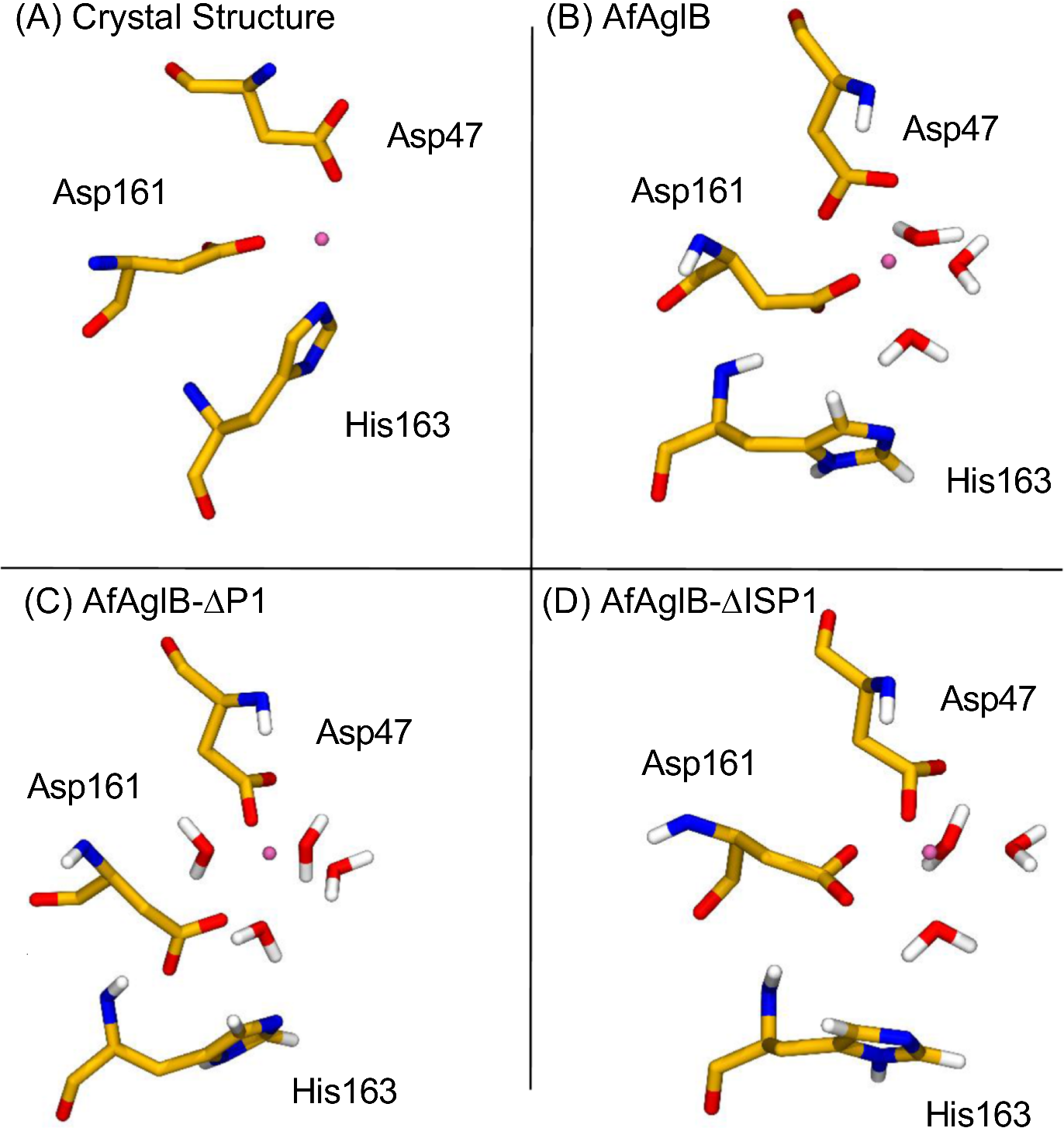
Snapshots of representative coordination geometries for the (A) crystallographic structure and each system studied: (B) AfAglB; (C) AfAglB-ΔP1; and (C) AfAglB-ΔISP1. Coordination numbers varies between 6 and 7, depending on the number of waters nearby or the denticity of the amino acids in the coordination complex. Residues from the catalytic site are represented in yellow licorice, while the rest of the protein, the membrane, and other water molecules are hidden for better visualization. Zn^2+^ ion is represented as a pink sphere.

**Table 1.**
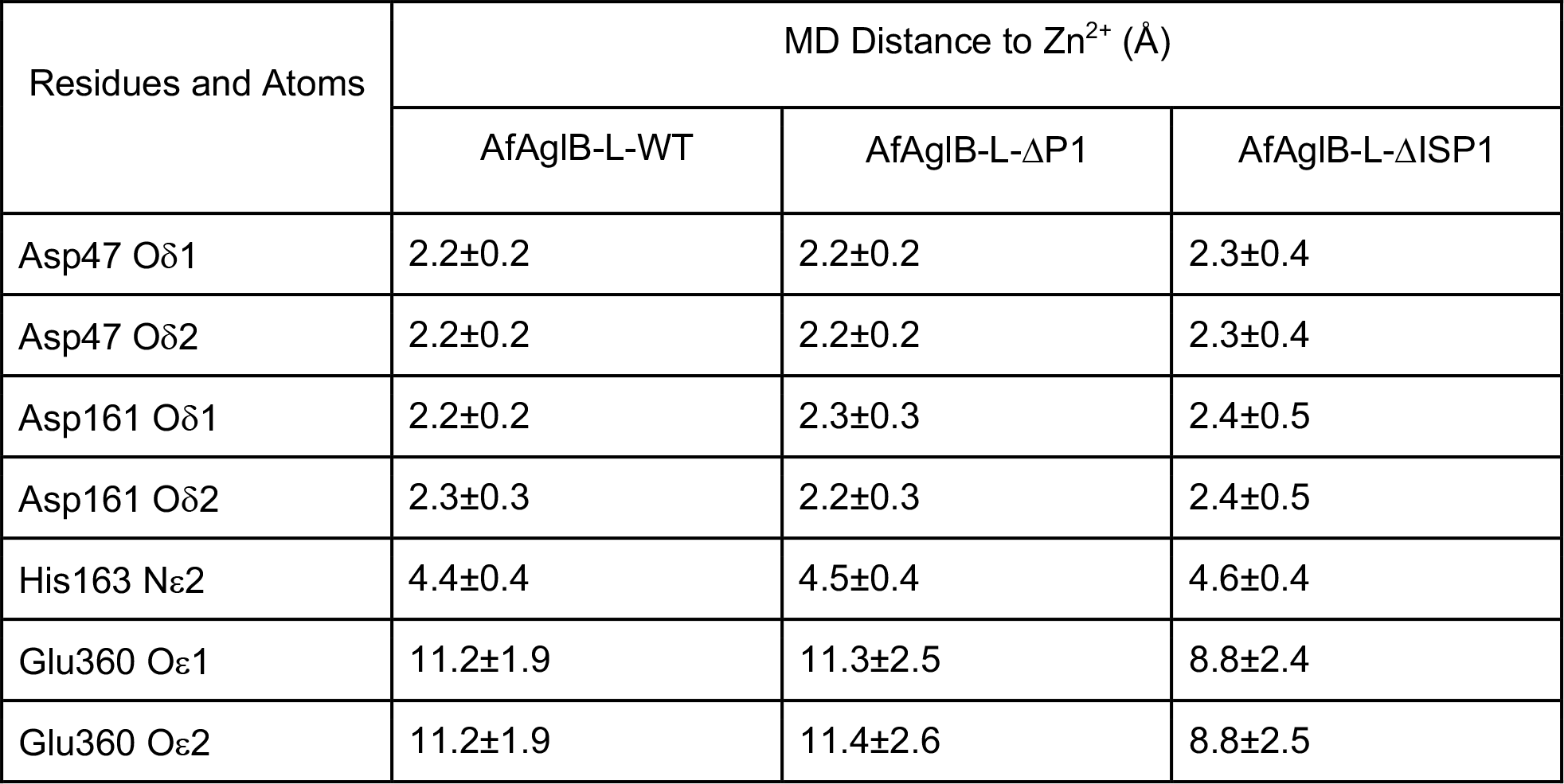
Distances from catalytic residues to Zn^2+^ ion during MD simulations (Å)

| Residues and Atoms | MD Distance to $Zn^{2+}$ (Å) | | |
| --- | --- | --- | --- |
| | AfAgIB-L-WT | AfAgIB-L-ΔP1 | AfAgIB-L-ΔISP1 |
| Asp47 Oδ1 | 2.2±0.2 | 2.2±0.2 | 2.3±0.4 |
| Asp47 Oδ2 | 2.2±0.2 | 2.2±0.2 | 2.3±0.4 |
| Asp161 Oδ1 | 2.2±0.2 | 2.3±0.3 | 2.4±0.5 |
| Asp161 Oδ2 | 2.3±0.3 | 2.2±0.3 | 2.4±0.5 |
| His163 Nε2 | 4.4±0.4 | 4.5±0.4 | 4.6±0.4 |
| Glu360 Oε1 | 11.2±1.9 | 11.3±2.5 | 8.8±2.4 |
| Glu360 Oε2 | 11.2±1.9 | 11.4±2.6 | 8.8±2.5 |

## Discussion

The deletion of structural units from the C-terminal domain of AfAglB-L impacts on many aspects of the enzyme. The TM domain appears to be the least influenced region, as we expected, since it is far from the mutations area, besides being embedded within the lipid environment, which provides stabilizing interactions. As seen by analyzing the structural fluctuations and divergence to the crystal structure, the CC unit of AfAglB-L-ΔISP1 was the most affected by the deletions, showing larger fluctuations throughout this unit, including a modification in the behavior of the kinked helix that carries the DK motif and in the WWDXGX motif area. As tested by previous reports^12,40^, this modification could negatively influence sequon recognition by the OST, rendering a poor protein binding and, ultimately, reducing the enzyme activity. AfAglB-L-ΔP1 is also influenced by the CC unit, but to a lesser extent; most of its destabilization is observed in the TH-Bundle, interfaced with the IS unit. The increased flexibility of both of these areas may pose as a drawback because not only it modifies the protein binding site interface with the acceptor protein, but it might also propagate to the CC unit, further destabilizing it in other areas, including the DK motif and the area near the LLO cavity. The presence of P1 in AfAglB-L-WT stabilizes the IS unit by the interface between the two units, interacting with helix aC of the TH-Bundle and the loop connecting it to the CC unit.

PCA technique is reported as a valuable tool to assess conformational sampling and convergence of simulations^41-43^. In our work, by using PCA we identified important aspects involving the conformational ensembles of the studied systems: 1) In spite of none of the systems reaching convergence on the PCA metric during our simulations, AfAglB-L-WT was the most dynamically stable system, since it was able to achieve stable energy basins, sample this areas, and had more overlap between its replicates, as well as a smaller drift along the PCs. The mutations affected the behavior of AfAglB-L, leading to huge drifts along the conformational space with just a few clearly defined basins being explored. 2) When evaluated in the same subspace, it is evident that the double unit deletion has a heavier impact on protein structure than the single deletion. In this case, none of the domains from AfAglB-L-ΔISP1 demonstrated a stable behavior, keeping the same projection patterns observed for the whole protein. AfAglB-L-WT and AfAglB-L-ΔP1, contraringly, exhibited some superimposition, which is probably due to the dominance of AfAglB-L-ΔISP1 largely fluctuating motions. Individual analyses for AfAglB-L-WT and AfAglB-L-ΔP1 systems confirmed that the overlap between their replicates is low, as well as highlighted the instabilities in the conformational profile of AfAglB-L-ΔP1 when compared to WT. Therefore, we conclude that the mutated systems are mostly unable to keep a similar conformational profile to AfAglB-L-WT, possibly interfering with the enzyme action, since the specific conformations that allow the binding of substrates may not be available on these systems.

Coordination of the metal ion at the binding site was very stable, indicating that, despite all the disturbance in the protein structure caused by the deletions, the enzyme could still be catalytic competent at some level, as long as the two substrates manage to bind in the cavities. A puzzling aspect of the coordination at the catalytic site involves His163. This residue is coordinating the metal ion in the crystal structure, displaying a distance of 2 Å from the nitrogen atom Nε2 to Zn^2+^. It is part of the DXD motif (DXH in AfAglB-L) of glycosyltransferases, equivalent to Asp156 in ClPglB OST, which has not been identified as one of the coordinating residues in its crystallographic structure^7^. These structures were obtained in different states, the apo (AfAglB-L) and the cocrystallized with the peptide acceptor (ClPglB), therefore one could assume that Asp156 and His163 may have a coordinating role only when the protein is in its apo state. However, both in this work and in a previous report from ClPglB simulations^44^, we did not observe this coordination participation during our simulations. Instead, a bidentate coordination by Asp47 (Asp56 in ClPglB) and water molecules mediated this interaction, maintaining the octahedral coordination geometry. Differently, a recent work^45^ also involving MD simulations of PglB, observed a coordination participation from Asp156 in all states studied. This way, the role of these residues is not clear yet, despite being widespread in this enzyme family^46^, as well as being fundamental to the enzyme activity^47^. One possibility is that His163 (and Asp156 in ClPglB) serves as a “backup” coordination ligand, preventing the metal ion from escaping the catalytic site by bridging water molecules interactions or by acting in situations where the protein substrate is absent. Another possibility would be that this residue is involved in catalysis by supporting the binding or the stabilization of the LLO substrate, since the monosaccharide at the reducing-end of the glycan is a glucose with three sulfates in positions C2, C3 and C6 and could be accommodated in a pocket consisting of the conserved residues His81, His162, His163, and Trp215. Of course, we have to consider that MD simulations do not handle electronic properties directly, which could be a caveat that prevented us from observing a correct description of the His163 interactions. Glu360 never reached the metal ion during any of the simulations, hence, the binding site was never fully complete. This strengthens the hypothesis that acceptor protein binding is important for the formation of the complete binding site^7,9^.

The data obtained for the catalytic site coordination corroborates previous data from PfAglB, where the catalytic activity of PfAglB-ΔIS was not affected by the deletion^11^. Contrastingly, the global motions and the binding interface, as seen by the PCs motions on its extreme projections (Figure S2), disagree with these observations. In AfAglB-L-WT, motions are dominated by EL5 from the TM unit, and by a rearrangement of the IS unit, similarly to motions identified in ClPglB dynamics^44^, reinforcing the observations that the IS unit is intrinsically flexible. In AfAglB-L-ΔP1, there is a twisting motion between TM and PP domains, whereas, in AfAglB-L-ΔISP1, we identified huge collective motions, especially on the PP domain, disorganizing the protein binding site. Comparing the PP domain of the wild-type structures from both species (Figure 7), we identified possible explanations for these discrepancies: the IS unit from PfAglB is a stable β-barrel structure, and the deletion (residues 603-678) allows residues 602 and 679 to be easily connected, thus causing less perturbations in the enzyme core than our deletion in AfAglB-L-WT (which included the TH-Bundle and one conserved β-sheet at the enzyme core). Indeed, the deletion in PfAglB permits a structural organization of the CC unit that resemble the other two paralogs from *A. fulgidus*, AfAglB-S1 and AfAglB-S2, which do not portray IS units. These observations indicate that structural units from PfAglB could act more independently when compared to AfAglB. Furthermore, PfAglB possesses two periphery units (P1 and P2), one that occupies the same place as P1 in AfAglB-L-WT, and another that encircles the area above the LLO cavity. As seen in our RMSF plots (Figure 3), this area has many unstable regions with fluctuations that are largely increased as a side-effect of the deletions. Flexibility of the same nature and at the same regions was also seen in ClPglB MD simulations^44^, confirming the disordering of these areas. Possibly, the presence of the β-strand rich P2 unit provides additional stabilization for PfAglB, protecting CC unit from unfolding, as well as allowing the deletion of IS to be tolerable. Perhaps, if AfAglB-L-WT possessed both P1 and P2, it could also tolerate the deletion of IS, as the single deletion of P1 was less aggressive to the structural organization of the enzyme core than the double deletion. Besides, the large-amplitude motions observed after both deletions in AfAglB-L would be contained by the presence of P2, promoting diminished fluctuations at the PP domain.

**Figure 7.**
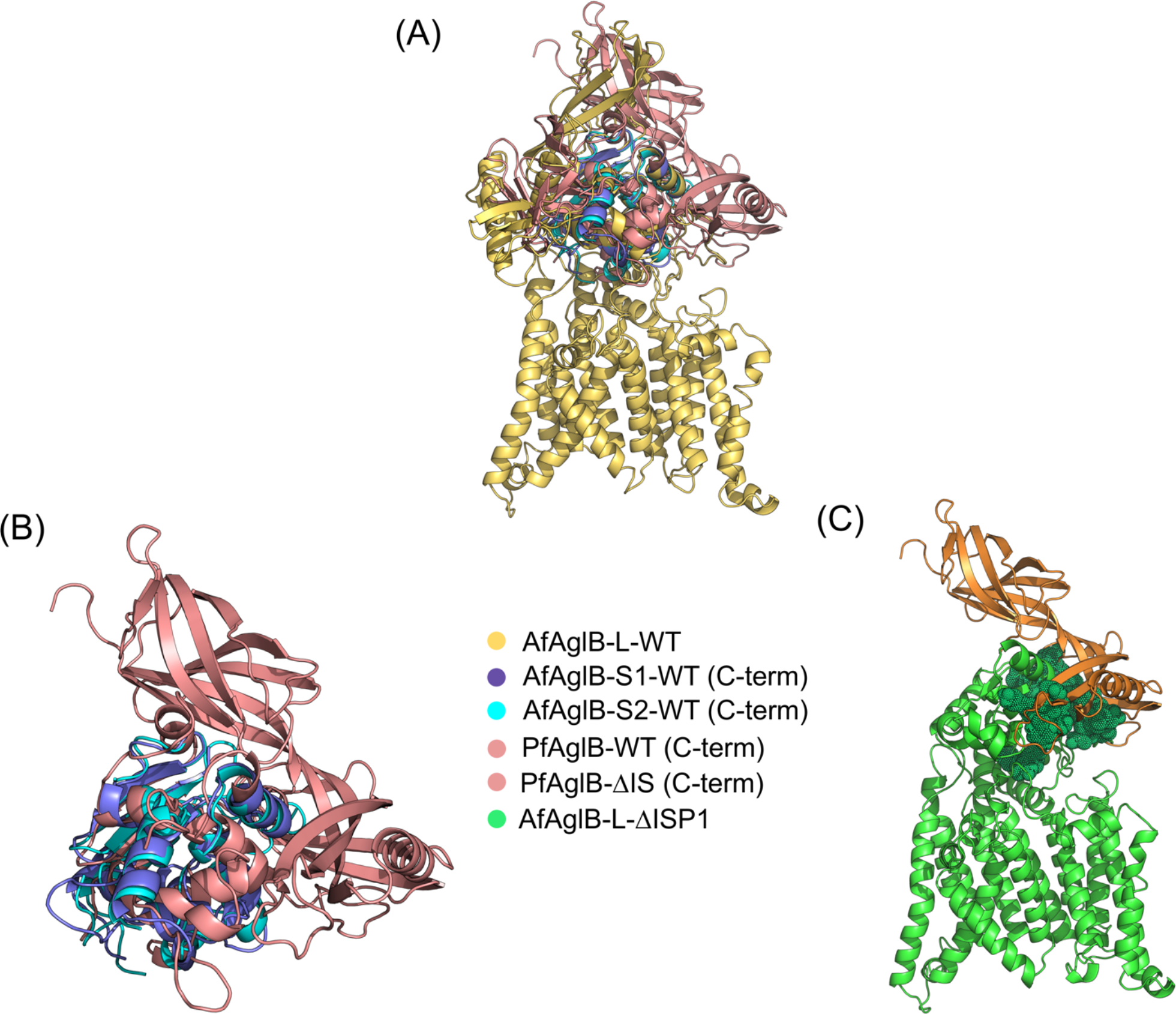
Structural comparison between AfAglB-L, AfAglB-S1 and S2, and PfAglB (PDB ID 2ZAI)^12^. (A) AfAglB-L with all other C-terminal domains aligned to its corresponding domain. (B) PfAglB with the deletion of IS unit folds in a way that resembles the other two paralogs from *A. fulgidus*. (C) Alignment of the C-terminal domain of PfAglB-WT to AfAglB-ΔISP1 left the P1 and P2 units (orange) aligned near a sensible area (green spheres) of the mutated system (AfAglB-ΔISP1), possibly protecting that region in a hypothetical chimeric AfAglB-L.

## Conclusions

Structural studies of OSTs have been constantly advancing, enabling a better understanding of the molecular basis involving N-glycosylation modifications, which, in turn, may provide new tools for the glycoengineering of biomolecules. Also, many experiments have been performed regarding these enzymes catalytic mechanism, usually evaluating residues from CC unit and TM domain, as well as studies designed to enhance their activities. However, to this date, only one previous work has assessed the role of other structural units from OSTs^11^. In this work, we try to contribute for the structural comprehension of OSTs and increase the knowledge about structural units dynamics and function. By employing MD simulations, we could verify that both P1 and IS units provide fundamental stabilizing interactions that permits both structural integrity of AfAglB-L in extreme environments, such as the high temperatures faced by *A. fulgidus*, along with the preservation of the enzyme conformational profile and binding interface, important features for its proper functioning. Combining our findings with previous data from literature, we identified that there might be some interchangeability between structural units. For example, one could transfer a P1 unit from AfAglB or P1 and P2 from PfAglB to another OST, aiming to achieve a new class of engineered chimeric thermostable OSTs. PglB from the *Campylobacter* or *Desulfovibrio* genera would be good candidates for these approaches, since they are found in mesophilic species and have a well-known promiscuity^20^. We hope that this work may support the development of more assessments regarding OSTs structural units.

## Acknowledgments

Research was supported by the Centro Nacional de Supercomputação of the Universidade Federal do Rio Grande do Sul (CESUP/UFRGS), CNPq (Conselho Nacional de Desenvolvimento Científico e Tecnológico), CAPES (Coordenadoria de Aperfeiçoamento de Pessoal de Nivel Superior), and FAPERGS (Fundação de Amparo à Pesquisa do Estado do Rio Grande do Sul).

