## Supporting Information File for "Assessment of Structural Units Deletions in the Archaeal Oligosaccharyltransferase AglB"

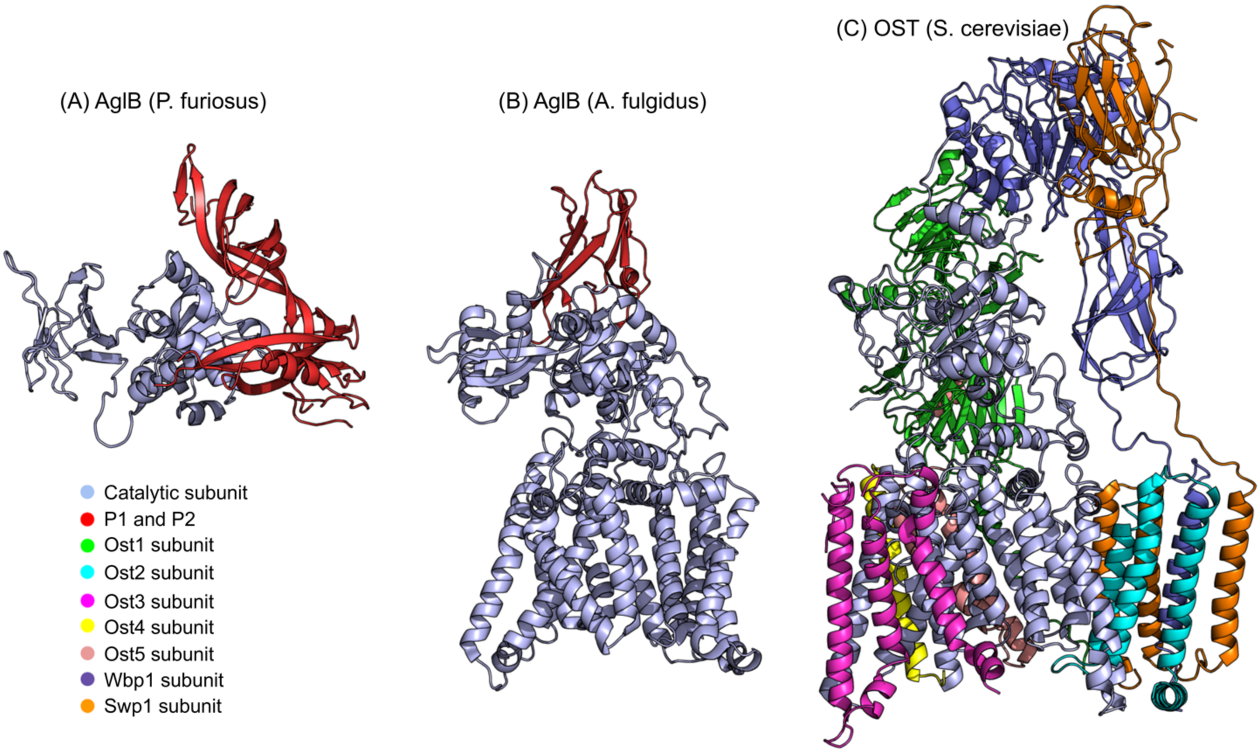


**Figure S1.** OST structural comparison of units and subunits on the archaea and eukarya domains. (A) AglB from *Pyrococcus furiosus* species with both P1 and P2 structural units (C-terminal domain only) (PDB ID: 2ZAI). (B) AglB from *Archaeoglobus fulgidus* with a P1 structural unit. (C) OST complex from *Saccharomyces cerevisiae* possessing eight subunits (PDB ID: 6C26). Despite being located in similar areas, P1, P2, and Wbp1 demonstrate very distinct organization and foldings between them.

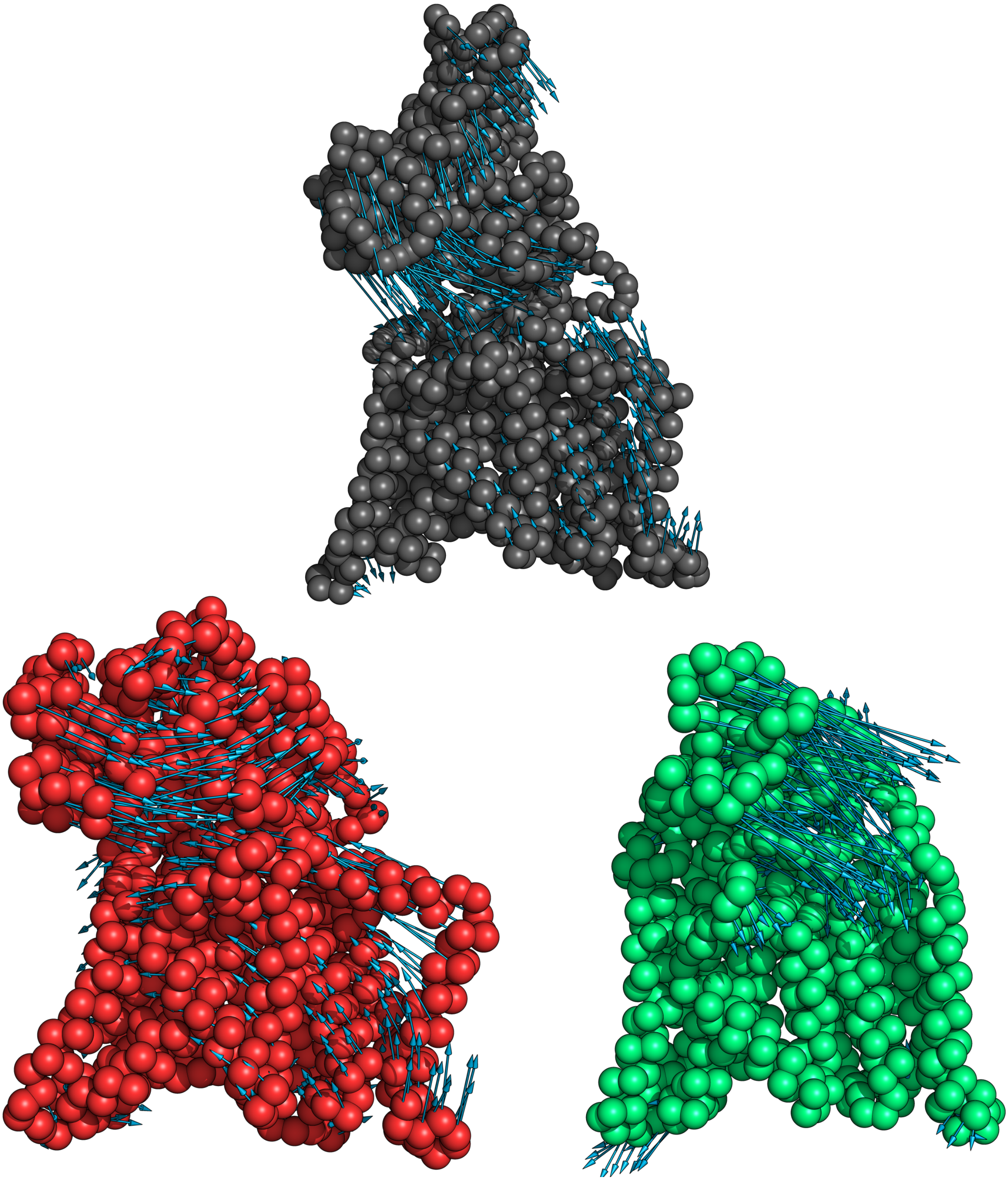

**Figure S2.** Porcupine plots depicting the movement described by the extreme projections of principal component 1 (PC1) of each group of replicates for the studied systems. AfAglB-L is represented by black spheres, AfAglB-L-ΔP1 is represented by red spheres, and AfAglB-L-ΔISP1 is represented by green spheres. Blue arrows indicate the direction of the motion of each protein area.
